# Systematic genetic interaction studies identify histone demethylase Utx as potential target for ameliorating Huntington’s disease

**DOI:** 10.1101/228486

**Authors:** Wan Song, Nóra Zsindely, Anikó Faragó, J. Lawrence Marsh, László Bodai

## Abstract

Huntington’s Disease (HD) is a dominantly inherited neurodegenerative disease caused by alterations in the huntingtin gene (*htt*). Transcriptional dysregulation is an early event in HD progression. Protein acetylation and methylation particularly on histones regulates chromatin structure thereby preventing or facilitating transcription. Although protein acetylation has been found to affect HD symptoms, little is known about the potential role of protein methylation in HD pathology. In recent years, a series of proteins have been described that are responsible for methylating and demethylating histones as well as other proteins. We carried out systematic genetic interaction studies testing lysine and arginine methylases and demethylases in a *Drosophila melanogaster* HD model. We found that modulating methylation enzymes that typically affect histone positions H3K4, H3K36 or H3K79 had varying effects on HD pathology while modulating ones that typically affect constitutive heterochromatin marks at H3K9 and H4K20 generally had limited impact on HD pathology. In contrast, modulating enzymes acting on the facultative heterochromatin mark at H3K27 had specific effects on HD pathology, with reduction of the demethylase Utx rescuing HTT induced pathology while reducing PRC2 complex core methylase components led to more aggressive pathology. Further exploration of the mechanism underlying the methylation-specific interactions suggest that these lysine and arginine methylases and demethylases are likely exerting their influence through non-histone targets. These results highlight a novel therapeutic approach for HD in the form of Utx inhibition.

## INTRODUCTION

Huntington’s disease (HD) is a fatal, neurodegenerative disorder caused by a CAG repeat expansion in the huntingtin (*htt*) gene, which encodes an abnormally long polyglutamine (polyQ) repeat in the HTT protein (1). In spite of its monogenic nature, the pathogenesis of HD is complex. Prior studies have identified over 700 proteins interacting with mutant HTT, which are involved in transcription, DNA maintenance, cell cycle regulation, cellular organization, protein transport, energy metabolism, cell signaling, and protein homeostasis (2).

Multiple studies have noted that altered modification of lysine residues is associated with HD (3-5). This coupled with observed transcription changes in HD has led to the idea that dysregulated lysine modifications might affect pathogenesis by altering histone modifications and thus transcription. Acetylation of histones is a well-known event that affects gene expression by relaxing chromatin structure. Protein methylation occurs on lysine and arginine residues and in many cases affects the same lysine residues as acetylation suggesting a potential interplay between their effects on cellular and pathogenic processes. Consistent with this, several studies have identified altered methylation patterns in HD patients and models (6-8).

However, histones are only one of many substrates targeted by lysine modifying enzymes. Both acetylation and methylation affect a wide-array of proteins (for reviews see (9, 10)) including histones and non-histone proteins with a wide-variety of functions, including transcriptional regulators, signaling molecules, metabolic enzymes and components of the cytoskeleton. Furthermore, the sets of enzymes participating in the modifications of histones and other proteins are not exclusive but overlapping. For example, the tumor suppressor p53 can be acetylated or methylated at several lysine or arginine residues, to influence its stability and activity (11-13). The lysine methyltransferases Set9 (12) and Set8/PR-Set7 (14) that act on p53 also modify histone H3 at lysine K4 (15) and histone H4 at lysine K20 (16), respectively. Thus, specific modifying enzymes can affect transcription by modifying both histone and non-histone proteins.

Despite the high number of non-histone acetylation and methylation targets, the majority of studies investigating the role of post-translational modifications (PTMs) on the pathogenesis of Huntington's disease have focused on the regulation of gene expression by altered modifications of histones. Transcriptional control relies on chromatin structure, which is regulated in part by PTMs of unstructured N-terminal regions of histones that directly change the intrinsic biophysical properties of local chromatin or indirectly allow or prevent docking of effector molecules (17-19). Histone methylation status determines the chromatin environment and thereby regulates transcription (20). Histone methylation marks can be divided into two major categories: active or repressive chromatin marks (21, 22) (Table 1). Simplistically, methylated H3K4, H3K36 and H3K79 are associated with actively transcribed genes (23, 24), whereas methylated H3K9, H4K20 and H3K27 are usually associated with inactive regions (25). *Drosophila melanogaster* has single orthologs for most of human histone methylase and demethylase genes and can be an ideal *in vivo* system to test the role of various histone methylation modifying enzymes on Huntingtin pathology.

**Table 1.** A summary of fly methylases and demethylases tested in the study. The known histone site targets, CG number and human homologs were listed. Based on their general genomic function, the genes were grouped to transcriptionally active or repressive marks, which were further categorized into marks associated with constitutive or facultative heterochromatin marks. The protein arginine methyltransferases tested were also included.

| Site | Function | Fruit fly protein | CG identifier | Human homolog |
| --- | --- | --- | --- | --- |
| ***Transcriptionally Active Chromatin*** |  |  |  |  |
| **H3K4** | methylation | Set1 | Cg40351 | SETD1A, SETD1B |
|  |  | Trx | Cg8651 | MLL1/2 |
|  |  | Trr | Cg3848 | MLL3/4 |
|  |  | not found |  | SET7/9 |
|  | demethylation | Su(var)3-3 | Cg17149 | LSD1 |
|  |  | Kdm2 | Cg11033 | KDM2B |
|  |  | Lid | Cg9088 | JARID1 |
| **H3K36** | methylation | Set2 | Cg1716 | SET2 |
|  |  | Ash1 | Cg8887 | ASH1 |
|  |  | NSD | Cg4976 | NSD1 |
|  |  | not found |  | SMYD2 |
|  | demethylation | Kdm4A | Cg15835 | JMJD2 |
|  |  | Kdm2 | Cg11033 | JHDM1A/1B |
|  |  | Kdm4B | Cg33182 | JMJD2 |
| **H3K79** | methylation | Gpp | Cg42803/Cg10272 | DOT1L |
|  | demethylation | not found |  | not found |
| ***Transcriptionally Inactive Chromatin*** |  |  |  |  |
| ***Constitutive heterochromatin marks*** |  |  |  |  |
| **H3K9** | methylation | Egg | Cg12196 | ESET |
|  |  | Su(var)3-9 | Cg43664 | SUV38H |
|  |  | G9a | Cg2995 | G9a |
|  | demethylation | JHDM2 | Cg8165 | JHDM2 |
|  |  | Kdm4B | Cg33182 | JMJD2 |
|  |  | Kdm4A | Cg15835 | JMJD2 |
| **H4K20** | methylation | PR-Set7 | Cg3307 | SET8 |
|  |  | Hmt4-20 | Cg13363 | SUV420H |
|  | demethylation | not found |  | PHF8 |
| ***Facultative heterochromatin marks*** |  |  |  |  |
| **H3K27** | methylation | E(z) | Cg6502 | Ezh2 |
|  | demethylation | Utx | Cg5640 | UTX/UTY, JMJD3 |
| ***Protein arginine methyltransferases*** |  |  |  |  |
| **unknown** | methylation | Art3 | CG6563 | PRMT3 |
| **unknown** | methylation | FBXO11 | CG9461 | PRMT9/FBXO11 |

In this study, we have sought to identify which methylation modifying loci affect HD pathology by systematically evaluating the effect of identified methyltransferases and demethylases in a *Drosophila* model of HD. In addition, we have sought to test the assumption of whether the key loci exert their effects by modifying histone residues or whether they exert their effect through other targets of methylation.

## RESULTS

### Modulating active chromatin histone methylation marks has varying effects on HD pathology

We first investigated methyltransferase and demethylase enzymes that act on lysine residues of histone tails whose methylation is associated with active transcription: H3K4, H3K36 or H3K79. H3K4 methylation represents a specific mark for epigenetic transcriptional activation and tri- and di-methylation on histone H3 at Lys 4 (H3K4me3/me2) is often found at active genes. H3K4me3 is highly enriched around transcriptional start sites (TSS) (26), while H3K4me2 is present both at TSS and coding regions of transcribed genes (26-31). Di- and tri-methylated forms of H3K36 residue are enriched at coding regions of actively transcribed genes and are involved in regulating transcriptional elongation (26, 32). H3K79 monomethylation is also enriched in the coding region of active genes, while the dimethylated form can be found on both the TSS and gene body of active genes (26).

In Drosophila, three non-redundant proteins - SET domain containing 1 (Set1), Trithorax (Trx) and Trithorax-related (Trr) - are found in complexes similar to the yeast COMPASS methyltransferase complex that are capable of methylating the H3K4 residues (33). Among these, Set1 was shown to act as the main global H3K4 di- and trimethylase throughout *Drosophila* development (34).

To determine whether H3K4 specific methyltransferases affect HD pathology, we conducted genetic interaction crosses to test the effects of reduced levels of methyltransferases (either by heterozygous loss of function mutations or by RNAi depletion) on HD phenotypes induced by the neuronal expression of human Httex1p-Q93 transgenes. When expressing mutant Httex1p-Q93 in all neurons with the pan-neuronal elav-GAL4 driver, adult flies exhibit reduced eclosion rates and neuronal degeneration as observed by decreased number of rhabdomeres in ommatidia in the pseudopupil assay (35). RNAi knockdown of *Set1* (GD12893) (33) in Httex1p-Q93 expressing flies did not influence viability (1.06 relative eclosion compared to control flies, not significant (N.S.), Fig. 1A) or cause a significant change in the number of rhabdomeres per ommatidium (+0.02 rhabdomeres per ommatidium in *Htt Set1*- RNAi/+ vs. *Htt* +/+ control, N.S., Fig. 1B), indicating that reducing Set1 has no impact on HD pathology.

**Figure 1.**
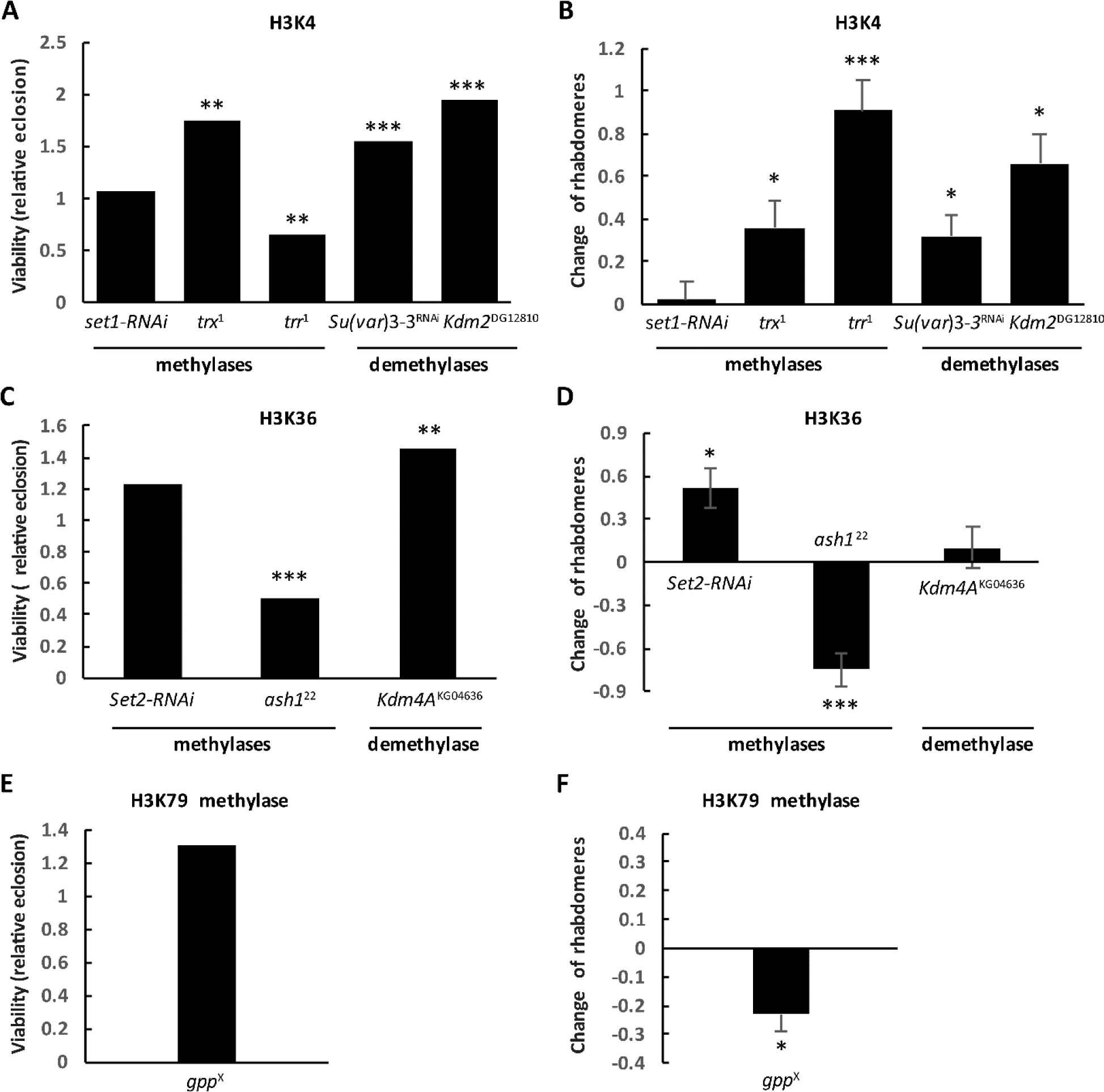
Methylases and demethylases acting on active marks H3K4 (**A** and **B**), H3K36 (**C** and **D**) and H3K79 (**E** and **F**) had varying effect on HD pathology. The graphs on the left show the effects of heterozygous loss of (**A**) H3K4, (**C**) H3K36, or (**E**) H3K79 specific methyltransferases and demethylases on Httex1p-Q93 induced reduced viability. Bars show relative eclosion calculated as the following ratio of eclosed progeny: ((Htt-expressing methylation mutants) / (Htt-expressing controls)) / ((Htt-non-expressing methylation mutants) / (Htt-non-expressing controls)). The graphs on the right show the effects of heterozygous loss of (**B**) H3K4, (**D**) H3K36, or (**F**) H3K79 specific methyltransferases and demethylases on Httex1p-Q93 induced neurodegeneration. Bars show the differences of average number of rhabdomeres per ommatidium in the eyes of methylation mutant Htt expressing flies and Htt expressing control siblings. Error bars represent standard error of mean. Significant differences are marked by *: P<0.05, **: P<0.01, ***: P<0.001.

In contrast, reducing Trx using the hypomorphic *trx^1^* allele (36), resulted in significant amelioration of HD pathology in Httex1p-Q93 expressing flies heterozygous for *trx^1^*. This was apparent both by a positive change in the number of rhabdomeres per ommatidium (+0.36 in *Htt trx^1^*/+ vs. *Htt* +/+, P=0.02, Fig. 1B) and significantly improved viability (relative eclosion of 1.75, P=0.003, Fig. 1A), indicating that Trx inhibition has a positive impact on HD pathology.

Reducing the levels of Trr by heterozygosity of the *trr^1^* null allele (37) also rescued the pseudopupil phenotype significantly (+0.91 rhabdomeres per ommatidium in *Htt trr^1^*/+ vs. *Htt* +/+ control, P=4×10^-5^, Fig. 1B), but resulted in reduced viability (0.64 relative eclosion compared to control, P=0.004, Fig. 1A). Similar results were also obtained by reduction of Trr in HD flies by heterozygous *trr^3^* which removes the SET domain (38) (Fig. S1 A and B). We conclude that unlike the globally acting Set1, reducing the levels of the more specialized H3K4 methylases, Trx and Trr, has a positive impact on Htt induced neurodegeneration.

Several enzymes are involved in the removal of methyl marks from H3K4 in *Drosophila*. Among them, Su(var)3-3/dLsd1 (*Drosophila* ortholog of lysine-specific demethylase 1) specifically demethylates H3K4me2 and H3K4me1 residues (39, 40) while Lid (little imaginal discs), the ortholog of the Jarid1 family of JmjC domain-containing proteins, demethylates H3K4me3 (41-43). In addition to Su(var)3-3 and Lid, Drosophila Kdm2 (Lysine (K)-specific demethylase 2) has also been shown to possess H3K4me3-specific lysine demethylase activity in adult flies (44). We have previously reported that reducing *Lid* ameliorates HD pathology (7). Similarly, an RNAi (GD9296) line of *Su*(*var*)3-3 rescued the pseudopupil phenotype (+0.32 rhabdomeres per ommatidium in *Htt Su(var)3*-*3*-*RNAi* vs. *Htt* +/+, P=0.03, Fig. 1B) and viability (1.54 relative eclosion, P=0.0001, Fig. 1A). Heterozygous loss of *Kdm2* by the amorphic *Kdm2^DG12810^* allele (45) has also ameliorated neurodegeneration (+0.66 rhabdomeres per ommatidium in *Htt Kdm2*/+ vs. *Htt* +/+, P=0.01, Fig. 1B) and improved viability (1.94 relative eclosion, P=2×10^-7^, Fig. 1A). Thus, reduction of H3K4 demethylases can have a positive impact on HD pathology.

Next we investigated the effects of modulating H3K36 methylation status on HD pathology. In *Drosophila,* H3K36 trimethylation is catalyzed by the product of the *Set domain containing 2* gene (Set2, dHypb) and dimethylation is mediated by the product of the *absent*, *small*, *or homeotic discs 1* gene (Ash1) (46) and *Mesoderm-expressed 4* gene (Mes4) (47). We tested an RNAi line of *Set2* (B24108) that was shown to eliminate H3K36 methylation in larvae (48). We saw significant rescue of HTT induced neurodegeneration when *Set2* was downregulated (+0.51 rhabdomeres per ommatidium in *Htt Set2*-*RNAi*/+ vs. *Htt* +/+, P=0.02, Fig. 1D) while no significant change in viability was observed (relative eclosion of 1.23, N.S., Fig. 1C).

The methyltransferase activity of Ash1 was initially reported to be directed towards H3K4, H3K9 and H4K20 (49, 50), however, recent evidence supports its dimethylation activity specifically towards H3K36 *in vitro* (46) and *in vivo* (51, 52). In contrast to the positive effect from reducing Set2, heterozygosity of *ash1^22^*, a null allele (53), resulted in an enhancement of HD pathology in Httex1p-Q93 expressing flies, apparent in both significantly more neurodegeneration (-0.75 rhabdomeres per ommatidium in *Htt ash1^22^*/+ vs. *Htt* +/+, P=0.0002, Fig. 1D) and a 50% reduction in relative eclosion rate (0.51, P=0.0001, Fig. 1C).

Demethylation of histone H3K36me3/me2 (and H3K9me3/me2) is mediated by the KDM4 subfamily, which belongs to a large family of Jumonji C (JmjC) domain-containing proteins (22, 43, 54). In *Drosophila*, Kdm4A is capable of demethylating H3K36me2 and H3K36me3 and regulates H3K36 methylation at multiple euchromatic sites (43, 55). To test the influence of Kdm4A on HD pathology we used the *Kdm4A^KG04636^* allele in which a P element inserted in the coding region of *Kdm4A* abrogates its expression (55). Reduced levels of Kdm4A (i.e. Htt-expressing flies that are also heterozygous for *Kdm4A^KG04636^*) led to significantly increased viability (1.45 relative eclosion, P=0.004, Fig. 1C) but had no effect on neurodegeneration (+0.10 change of rhabdomeres per ommatidium, N.S., Fig. 1D).

The *grappa* gene (*gpp*) is solely responsible for catalyzing the mono-, di-, and tri- methylation of H3K79 (56, 57) and has the unusual property of exhibiting phenotypes and genetic interactions that are characteristic of both Polycomb-group (PC-G) and Trithorax-group (TRX-G) genes (56). We tested two X-ray induced loss of function alleles of *gpp*, *gpp^X^* and *gpp^XXV^*. Heterozygous reduction of Gpp in Htt-expressing flies resulted in modest, yet significant increase in neurodegeneration (-0.23 rhabdomeres per ommatidium in *Htt gpp^X^*/+ vs. *Htt* +/+ control, P=0.01, Fig. 1F) with no significant impact on viability (relative eclosion of 1.31, N.S., Fig. 1E). Reducing Gpp by the other loss of function allele, *gpp^XXV^*, gave similar results with even stronger neurodegeneration (Fig. S1 A and B).

### Modulating histone marks specific for constitutive heterochromatin marks has limited impact on Htt pathology

Next we investigated the effects of enzymes modulating epigenetic marks specific for constitutive heterochromatin, H3K9 and H4K20 methylation, on HD pathogenesis.

H3K9 trimethylation represents a specific mark for epigenetic repression that recruits the Su(var)205/HP1 protein to methylated histones, which then induces heterochromatin formation (58-60). Three SET domain containing lysine methyltransferases are responsible for methylating the H3K9 residue in fruit flies: Eggless (Egg), the *Drosophila* ortholog of human SETDB1 and mouse ESET (61-63); Su(var)3-9, the *Drosophila* ortholog of human Su(var)3-9 (64); and G9a, the *Drosophila* ortholog of human and mouse G9a (65, 66). These enzymes have different expression patterns, preferred methylation substrates and functions (67).

Mutants or transgenic insertion lines of each of the three genes were tested for effects on HTT-mediated degeneration. Heterozygous reduction using a loss of function allele of *eggless*, *egg^1473^* (68) did not have any significant effect on either viability (1.29 relative elosion, N.S. Fig. 2A) or neurodegeneration (-0.1 change of rhabdomeres per ommatidium, N.S., Fig. 2B). Testing with an independent loss of function allele *egg^235^* (68) gave similar results (Fig. S2 A and B).

**Figure 2.**
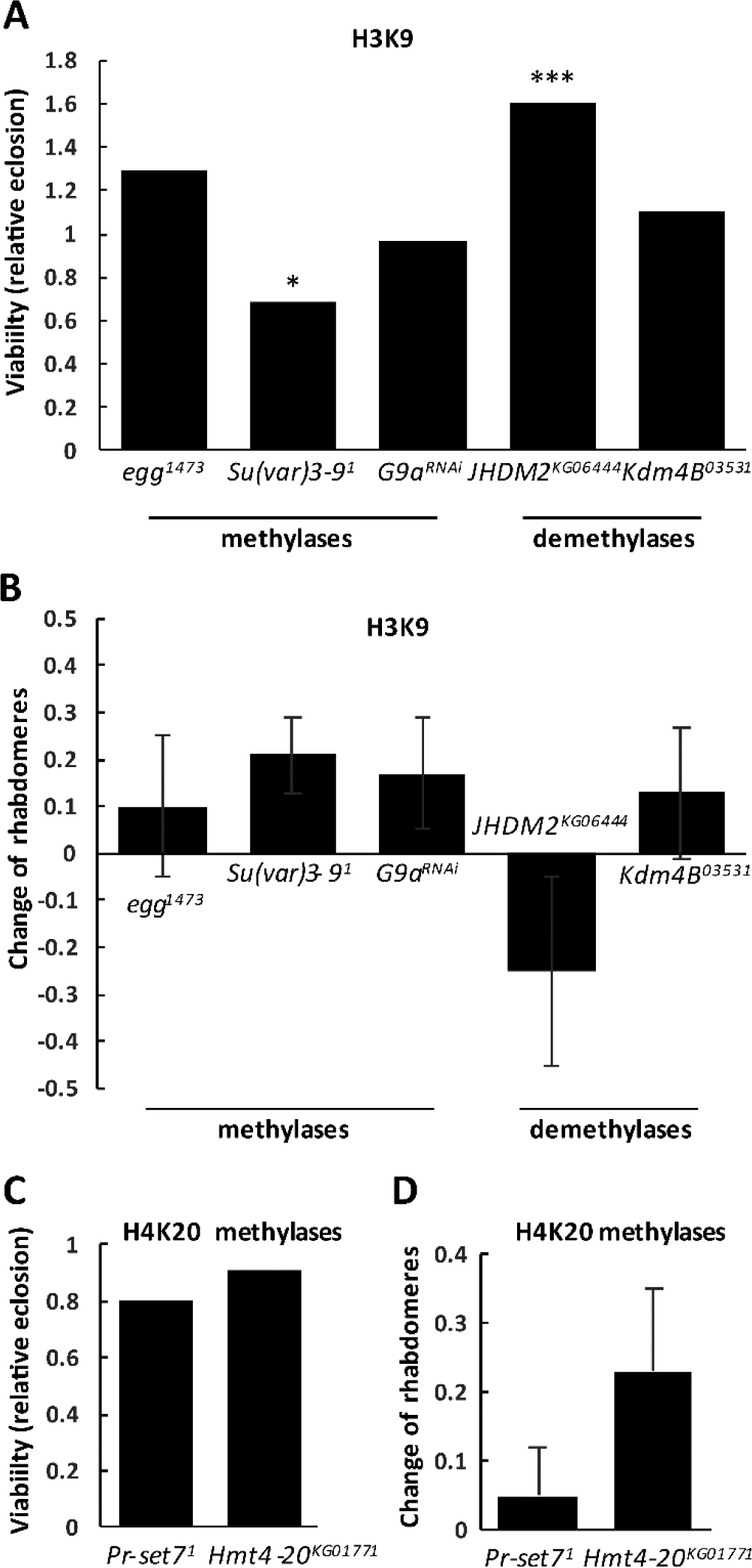
Modulating constitutive heterochromatin marks H3K9 and H4K20 methylation has no effect on HD pathology. (**A**) The effects of heterozygous loss of H3K9 specific methyltransferases and demethylases on Httex1p-Q93 induced reduced viability. Bars show relative eclosion calculated as the following ratio of eclosed progeny: ((Htt-expressing methylation mutants) / (Htt-expressing controls)) / ((Htt-non-expressing methylation mutants) / (Htt-non-expressing controls)). (**B**) The effects of heterozygous loss of H3K9 specific methyltransferases and demethylases on Httex1p-Q93 induced neurodegeneration. Bars show the differences of average number of rhabdomeres per ommatidium in the eyes of methylation mutant Htt expressing flies and Htt expressing control siblings, error bars represent standard error of mean. (**C**) The effects of heterozygous loss of H4K20 specific methyltransferases on Httex1p-Q93 induced reduced viability. Bars show relative eclosion. (**D**) The effects of heterozygous loss of H4K20 specific methyltransferases on Httex1p-Q93 induced neurodegeneration. Differences of average number of rhabdomeres per ommatidium in the eyes of methylation mutant Htt expressing flies and Htt expressing control siblings are shown. Error bars represent standard error of mean. Significant differences are marked by *: P<0.05, **: P<0.01, ***: P<0.001.

Heterozygous reduction of Su(var)3-9 with an amorphic allele *Su*(*var*)*3*-*9^1^* (69) resulted in no effect on neurodegeneration (+0.21 change of rhabdomeres per ommatidium, N.S., Fig. 2B) albeit reduced viability (relative eclosion of 0.69, P=0.02, Fig. 2A) in Htt-challenged flies. Testing with an independent amorphic allele, *Su*(*var*)*3*-*9^2^* (70), gave similar result on neurodegeneration but ameliorated viability (Fig. S2 A and B).

An RNAi line for *G9a*, GD9876, was confirmed by Q-RT-PCR to reduce G9a mRNA levels to 63% of control (Fig. S2 C). *Htt G9a*-*RNAi*/+ flies showed similar viability relative to control *Htt* +/+ (0.96 relative eclosion, N.S., Fig. 2A) and a non-significant change of rhabdomeres per ommatidium (+0.17, N.S., Fig. 2B). Heterozygous reduction using allele *G9a^MB11975^*, in which a Gal4 enhancer trap element is inserted into the 2nd exon of the gene, also did not significantly alter neurodegeneration but did reduce viability (Fig. S2 A and B).

The effect on HD pathology of reducing the activity of the H3K9 demethylases that exert the opposing activity was examined. JmjC-domain containing histone demethylase 2 (JHDM2), was shown previously to specifically demethylate H3K9 (71). Heterozygous reduction of *JHDM2* (*JHDM2^KG06444^*/+) did not affect neurodegeneration significantly (−0.25 change of rhabdomeres per ommatidium in Htt *JHDM2*/+ vs. *Htt* +/+, N.S., Fig. 2B) but improved viability (relative eclosion of 1.60, P=4×10^-5^, Fig. 2A) compared to the controls.

The demethylase encoded by the *Lysine* (*K*)-*specific demethylase 4B* (*Kdm4B*) gene was shown to act on both H3K9 and H3K36 residues (72). Using heterozygous *Kdm4B^03531^*, a P element insertion allele that results in substantial reduction in Kdm4B protein levels in heterozygotes (73), we did not observe a significant impact on either viability (relative eclosion rate of 1.10, N.S., Fig. 2A) or neurodegeneration (0.13 change of rhabdomeres per ommatidium, N.S., Fig. 2B).

Similar to H3K9 methylation, H4K20 mono- or tri-methylation represents a specific mark for epigenetic transcriptional repression. H4K20 trimethylation is mainly enriched in pericentric heterochromatin regions where it plays a central role in the establishment of constitutive heterochromatin (74). PR/SET domain containing protein 7 (PR-Set7) specifically acts to mono-methylate (75, 76) whereas histone methyltransferase 4-20 (Hmt4-20) is responsible for di- and tri-methylating H4K20 (77). We tested the effects of the hypomorphic *PR*-*Set1^1^* allele (78) on Htt induced pathology and did not see any significant impact on either viability (relative eclosion of 0.80 in *Htt PR*-*Set7^1^*/+ vs. *Htt* +/+, N.S., Fig. 2C) or neurodegeneration (+0.05 change of rhabdomeres per ommatidium, N.S., Fig. 2D). Similarly, the *Hmt4*-*20^KG01771^* allele, an allele with a P-element inserted in the 5’ UTR of the Hmt4-20-RA transcript, did not result in any significant difference in *Htt Hmt4*-*20*/+ flies compared to their control siblings (*Htt* +/+) in either viability (relative eclosion of 0.91, N.S., Fig. 2C) or change of rhabdomeres per ommatidium (-0.23, N.S., Fig. 2D).

Based on the results described above involving mutant Huntingtin and H3K9 methyltransferases and demethylases and H4K20 methyltransferases, we conclude that modulating constitutive heterochromatin marks does not exert a significant impact on HTT mediated HD pathology in Drosophila.

### Altered levels of enzymes modulating the facultative heterochromatin mark H3K27me3 impacts HD pathology

Tri-methylated lysine 27 on histone H3 (H3K27me3) is the most well-characterized mark for “facultative heterochromatin” and is critical for the repression of key transcriptional regulators during development (79). It was previously reported that full length Huntingtin acts as a facilitator of the multi-subunit histone H3 lysine 27 (H3K27) methyltransferase complex Polycomb Repressive Complex 2 (PRC2) in mice, and mouse embryos lacking Huntingtin display a series of phenotypes similar to that of PRC2-deficient embryos (80). We asked if modulation of enzymes acting on the H3K27me3 repressive mark influences expanded polyQ Huntingtin induced pathology. The Polycomb-group protein E(z) is the only protein found so far to be responsible for methylating H3K27 in *Drosophila* (81-83). Also, while there are multiple H3K27me3-specific demethylases, including UTX, UTY, and JmjD3 in mammalian cells, *Drosophila* have a single histone demethylase, Utx, capable of specifically demethylating di- and tri-methylated H3K27 (84). The fact that there is only one enzyme catalyzing each direction of methylation/demethylation makes testing H3K27 specific enzymes straightforward in Drosophila.

The null allele *E*(*z*)*^731^* has a nonsense mutation within the SET domain, the catalytic part responsible for the methyltransferase activity, and produces no detectable protein (82). Compared to Htt expressing control flies, Htt-expressing flies heterozygous for *E(z)^731^*/+ had significantly lower viability (relative eclosion of 0.54, P=3.0×10^-7^, Fig. 3A) as well as significantly fewer rhabdomeres per ommatidium (-0.34 less rhabdomeres per ommatidium compared to the control, P=0.02, Fig. 3B). Significant reduction in viability (79% relative eclosion rate, P=0.004) and neurodegeneration (-0.48 less rhabdomeres per ommatidium compared to control, P= 0.008) was also observed with another null allele, *E(z)^63^* (Fig. S3 A and B). These results indicate that reducing the E(z) methylase enhances HD pathology.

**Figure 3.**
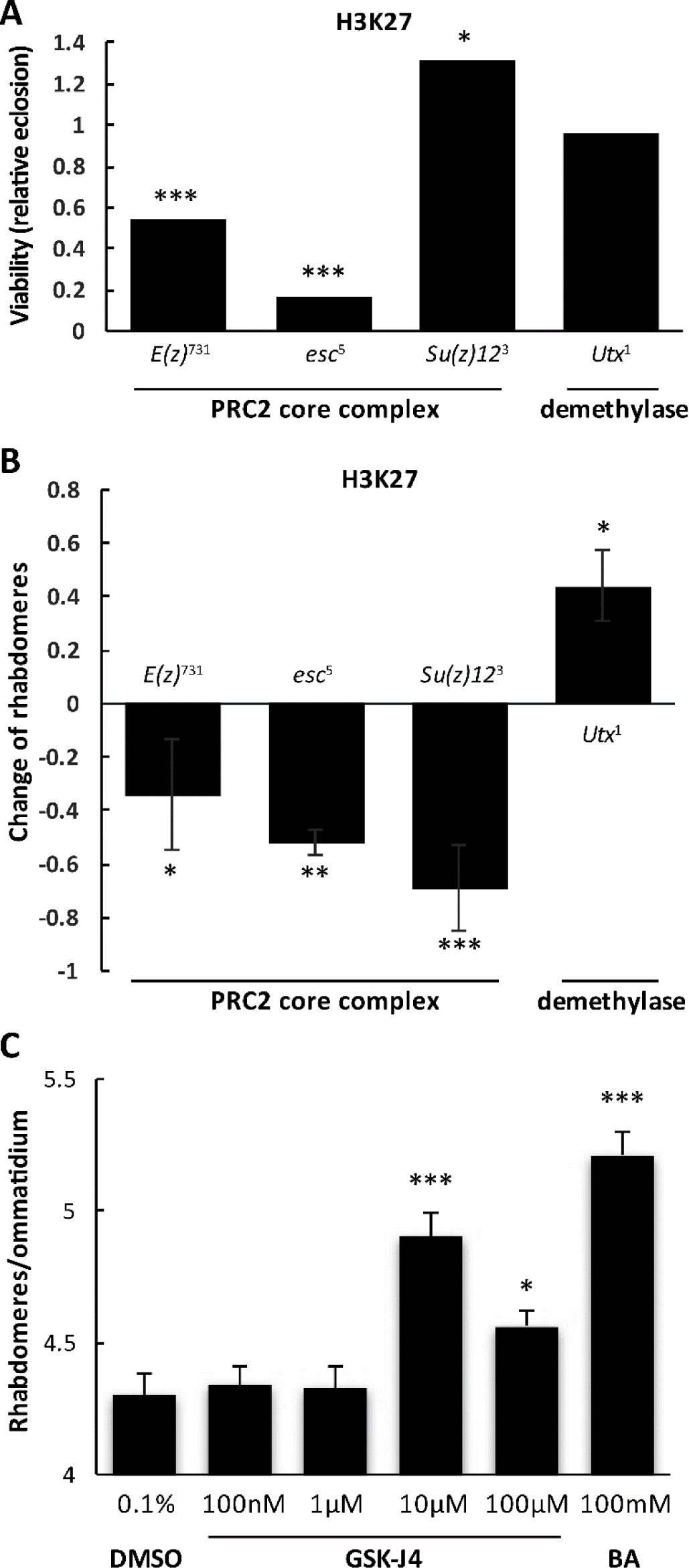
Modulating H3K27 methylation has specific effect on HD pathology. (**A**) The effects of heterozygous loss of subunits of the H3K27 specific PRC2 methyltransferase complex and Utx demethylase on Httex1p-Q93 induced reduced viability. Bars show relative eclosion calculated as the following ratio of eclosed progeny: ((Htt-expressing methylation mutants) / (Htt-expressing controls)) / ((Htt-non-expressing methylation mutants) / (Htt-non-expressing controls)). (**B**) The effects of heterozygous loss of PRC2 subunits and Utx on Httex1p-Q93 induced neurodegeneration. Bars show the differences of average number of rhabdomeres per ommatidium in the eyes of methylation mutant Htt expressing flies and Htt expressing control siblings. Error bars represent standard error of mean. (**C**) Utx inhibitor GSK-J4 is neuroprotective as shown by the significantly improved photoreceptor neuron survival at concentrations of 10 μM and 100 μM concentrations in flies challenged with Httex1p-Q93. No change in neurodegeneration was observed in the 0.1% DMSO vehicle control. 100 mM butyric acid served as positive control. Significant differences are marked by *: P<0.05, **: P<0.01, ***: P<0.001.

However, E(z) does not function as a single enzyme but as the catalytic subunit of the PRC2 protein complex (81), which besides E(z), contains three core subunits: Extra sex combs (Esc, human EED ortholog in *Drosophila*), Suppressor of zeste-12 (Su(z)12), and nucleosome-remodeling factor 55 (NURF-55) (81, 82). Esc is a WD repeat containing protein which binds specifically to trimethylated H3K27 whereas Su(z)12 is proposed to bind E(z) and promote PRC complex assembly (85). Both Esc and Su(z)12 are required for the integrity of PRC2 and PRC-mediated H3K27 methylation (86, 87). To investigate possible interactions of Htt with these components of the PRC2 complex, we carried out genetic interaction tests with mutant alleles of *Esc* and *Su(z)12* as well as non-core components of PRC2. Similar to E(z), heterozygous reduction of Esc by the *esc^5^* null allele enhanced pathology. We observed severely reduced viability (relative eclosion of 0.16, P=1.50x10^-12^, Fig. 3A) and significantly enhanced neurodegeneration (-0.52 less rhabdomeres per ommatidium in *Htt esc^5^*/+ vs. *Htt* +/+ flies, P=0.02, Fig. 3B). Heterozygous reduction of Su(z)12 by the loss of function allele *Su(z)12^3^* also resulted in significantly enhanced neurodegeneration (-0.69 less rhabdomeres in *Htt Su(z)12^3^*/+ than in *Htt* +/+, P=0.0007, Fig. 3B) albeit positive impact on viability (1.30 relative eclosion, P=0.03, Fig. 3A). In contrast, when we tested two non-core components of PRC2, Pcl and Jing, we did not observe any significant impact on HD pathology (Fig. S3 A and B). However, reduced levels of a third non-core PRC2 component, Jarid2, by heterozygous *Jarid2^MB00996^* null allele suppressed HD pathology as evidenced by significantly higher average number of rhabdomeres per ommatidium (+0.54 more than in controls, P=0.008, Fig. S3 B) and increased viability (3.18 relative eclosion rate, p=2.8×10^-15^, Fig. S3 A). As a comparison, we also tested Pc, a core component of PRC1, which also compacts chromatin but catalyzes H2A monoubiquitylation independently of PRC2 and H3K27me3 (88), and saw no significant effect on the survival of photoreceptor neurons (-0.23 less rhabdomeres per ommatidium in *Htt Pc*^1^/+ than *Htt* +/+, N.S., Fig. S3 B) albeit Htt *Pc*^1^/+ showed improved viability than in *Htt* +/+ control flies (relative eclosion of 1.99, P=0.003, Fig. S3 A).

Based on the above, we reasoned that reducing the activity of demethylase Utx that exerts an opposite enzymatic function on H3K27me3 might rescue HD pathology. First we tested the *Utx^1^* allele, which has a premature termination codon in the JmjC domain and is a strong hypomorph (89). Consistent with our predictions, heterozygous *Utx^1^* rescued the HD neurodegeneration phenotype, with the number of rhabdomeres per ommatidium significantly higher in *Htt Utx^1^*/+ flies compared with Htt-expressing control flies (+0.44 more rhabdomeres per ommatidium, P=0.02, Fig. 3B) while it had no significant impact on viability (relative eclosion of 0.96, N.S., Fig. 3A). Testing a second allele, *Utx^2^*, which contains a missense mutation in the C terminus resulting in a reduced amount of Utx protein (89), also gave similar results (Fig. S3 A and B). Therefore, contrary to the effect of reducing E(z), heterozygous reduction of Utx ameliorates mutant Huntingtin induced neurodegeneration. Thus, manipulations that tend to increase H3K27 methylation (e.g. lower Utx) tend to rescue HTT induced pathology while those that tend to decrease H3K27 methylation (e.g. reducing PRC2 core complex components) lead to more aggressive pathology.

Recently, a small molecule, GSK-J1, was developed to be a selective catalytic site inhibitor of H3K27me3-specific jumonji JMJ subfamily of proteins (90). Drosophila H3K27me3-specific demethylase Utx contains all the critical amino acids involved in GSK-J1 binding (data not shown). Since reducing Utx activity genetically rescued HD pathology, we explored the possibility of rescuing the HD pathology pharmacologically with this Utx inhibitor. We compared the effect of increasing concentrations of the H3K27me3-specific inhibitor, the cell-penetrating ester prodrug GSK-J4, butyrate (BA, positive control) and DMSO (negative control). As a histone deacetylase inhibitor shown in previous studies (35), BA is neuroprotective as Htt-expressing flies raised on 100 mM BA had significantly higher average number of rhabdomeres per ommatidium than DMSO treatment (5.21 vs. 4.3, P=4.05x10^-9^, Fig. 3C). Interestingly, flies on the higher concentrations (10 μM and 100 μM) of GSK-J4 showed higher number of rhabdomeres per ommatidium, with the average of 4.9 for 100 μM and 4.65 for 10 μM, which are both significant higher compared to the 0.1% DMSO control (P=2.00×10^-5^ and P=0.05, respectively. Fig. 3C). In contrast, flies on the lower concentrations (100 nM and 1 μM) of GSK-J4 showed no difference relative to the DMSO control (Fig. 3C). We also tested transferring the Htt-expressing flies to the drug-containing food after their eclosion but saw no difference between GSK-J4 and DMSO treatment (data not shown). Thus both genetically and pharmacologically, manipulation of the methylation activity of Utx holds promise as a novel therapeutic drug target for ameliorating HD symptoms.

### Partial loss of Utx reduces Htt protein aggregation

As genetic factors mediating H3K27 methylation or demethylation showed opposite effects on mutant HTT induced phenotypes suggesting a potentially attractive therapeutic target, we decided to further investigate the potential involvement of Utx in HD pathology. First, we needed to exclude the possibility that reduced Utx ameliorates HD phenotypes in the model because Htt transgene expression *per se* is affected by *Utx* mutations. We performed q-RT-PCR analysis on Htt-challenged *Utx* mutant (elav-GAL4/*w*; *Utx*^1^/+; UAS-Httex1p-Q93/+) or Htt-challenged control siblings (elav-GAL4/*w*; +/+; UAS-Httex1p-Q93/+) and found no significant difference in the level of Htt expression in *Utx^1^* heterozygotes (Fig. 4A). Similar results were also obtained with *Utx^2^* heterozygotes (Fig. S3 C). This indicates that the rescuing effect of Utx reduction on HD pathology is not due to a reduced level of Htt transgene expression.

**Figure 4.**
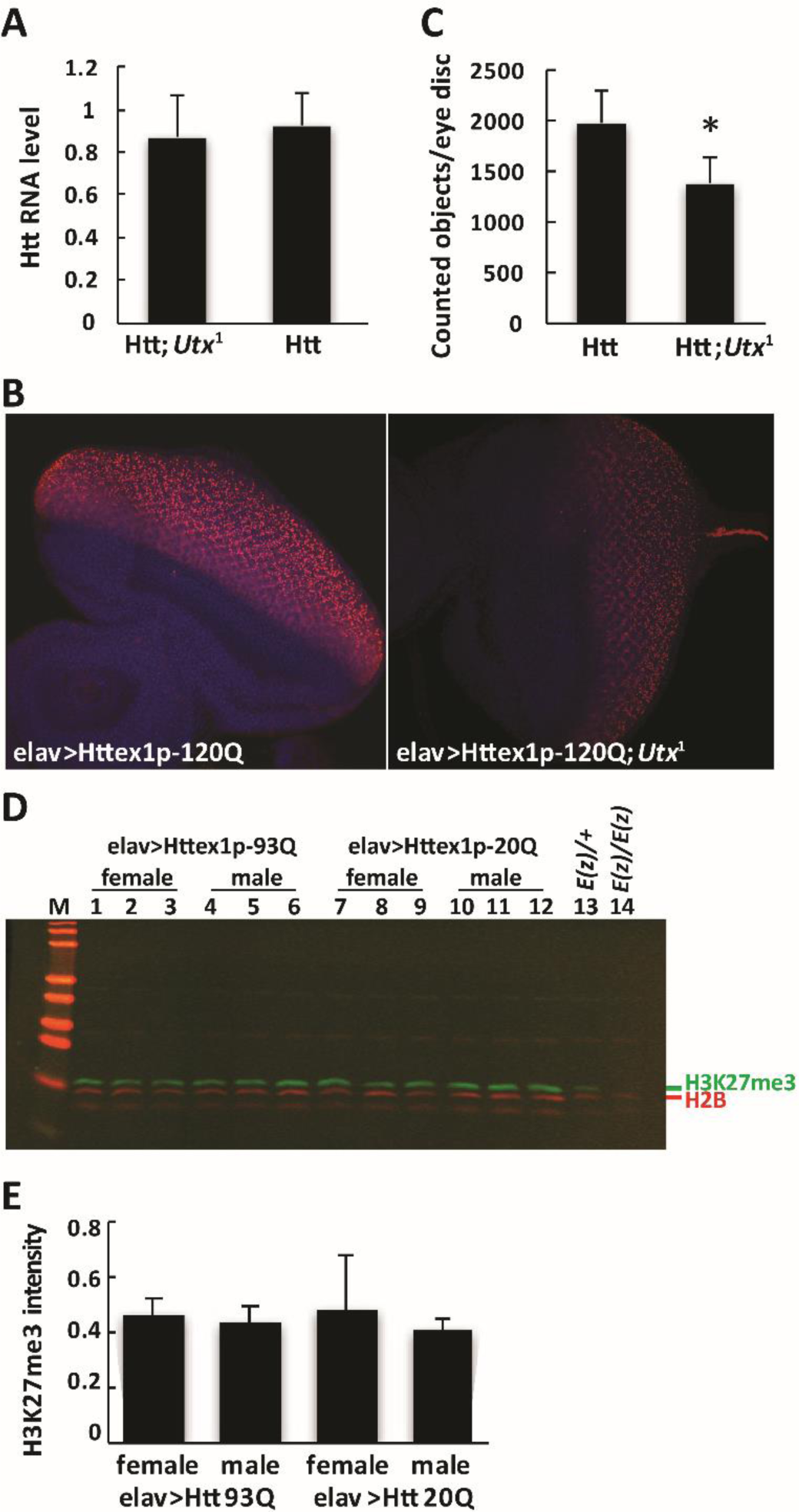
Reduction of *Utx* reduces Htt protein accumulation and aggregation but mutant Htt does not affect the methylation state of Utx targeted H3K27. (**A**) Heterozygous reduction of *Utx^1^* does not have a significant effect on Htt transgene expression as measured by Q-RT-PCR. (**B**) Htt protein accumulation and aggregation in eye discs of elav-GAL4/+; *Utx^1^*/+ Httex1p-Q120/+ larvae is reduced compared to control elav-GAL4/+; Httex1p-Q120/+ larvae. Representative compiled 3D confocal Z-stack images of fluorescent eye discs dissected from third instar larvae from indicated genotypes are shown. Tissues were stained with α-Htt antibody (1:2K VB3130, red) and Hoechst (blue). Images were taken at the same settings. (**C**) Quantification of α-Htt positive puncta in eye discs by 3D Object counting function of ImageJ (n>=4) shows that heterozygous reduction of *Utx* results in reduced Htt aggregation. (**D**) Representative quantitative Western blot showing similar signals from Htt challenged (elav-GAL4>Httex1p-Q93) or non-challenged (elav-GAL4>Httex1p-Q20) heads. Histones prepared from heads of females or males were probed on Western blots with antibody specific for trimethylated histone H3K27 (Millipore 07-449). Lanes 13 and 14 show diminished H3K27me3 signal from histones prepared from transheterozygotes (Lane 14, *E(z)^731^*/*E(z)^63^*) compared to heterozygotes (lane 13, *E(z)*/+). Three biological replicates were analyzed for females and males of each genotype. (**E**) Quantification of H3K27me3 signal intensities normalized to H2B signal from Western blots show similar levels in elav-GAL4>Httex1p-Q93 and elav-GAL4>Httex1p-Q20 male and female head samples.

As aggregation of mutant Htt is a hallmark feature of HD, we investigated whether reducing Utx affects HTT protein accumulation and aggregation. To test this, we drove the expression of mutant Huntingtin exon1 in neurons with elav-GAL4 in the presence or absence of *Utx* mutations and examined Htt protein accumulation and aggregation in larval eye discs. When the tissues were immuno-stained with Htt exon1-specific antibody (VB3130), we observed stronger Htt protein aggregation with the expression of Httex1p-Q120 compared to Httex1p-Q93, therefore we used Httex1p-Q120 for further immunofluorescence image analysis (Fig. 4B). By counting Htt aggregate-positive puncta on complied 3D images, we found that eye discs of Htt expressing flies heterozygous for *Utx* (elav-GAL4/+; *Utx^1^*/+; UAS-Httex1p-Q120/+) had significantly less HTT puncta than eye discs of control siblings (elav-GAL4/+; +/+; Httex1p-120Q/+) (Figure 4C). This result indicates that heterozygous reduction of *Utx* decreased the level of HTT protein aggregation.

### Interrogation of H3K27 methylation levels detect no change in HD flies

Our genetic interaction tests suggested that expression of mutant Htt might lead to decreased H3K27 methylation and reduction of Utx activity might have rescued neurodegeneration by correcting the histone methylation imbalance. To test this possibility, we first investigated whether global H3K27 trimethylation was affected by Htt expression. We quantified the level of H3K27me3 in the heads of flies expressing Httex1p-Q93 vs. Httex1p-Q20 controls by the elav-GAL4 driver using quantitative Western blot analysis (Fig. 4D and 4E). In contrast of the significantly diminished H3K27me3 levels in larval samples of *E(z)* null trans-heterozygotes *E(z)^731^*/*E(z)^63^* (Lane 15 in Fig. 4D) relative to *E(z)^731^/+* heterozygotes (Lane 14 in Fig. 4D), we did not observe any significant decrease in H3K27 trimethylation level in fly head samples from either sex expressing Httex1p-Q93 (Lane 2~7) compared to Httex1p-Q20 controls (Lane 8~13 in Fig. 4D, quantification in Fig. 4E), suggesting a lack of effect from HTT challenge on global level of H3K27me3 in HD flies.

Although we did not detect a global change in H3K27me3 levels in HTT expressing flies it is possible that HD challenge modifies the H3K27me3 marks on specific genes. We tested this by chromatin immunoprecipitation (ChIP). From the publically available modENCODE *Drosophila* adult head H3K27me3 ChIP-Chip or ChIP-seq datasets (91) (#372-male, ChIP-Chip; #346-female, ChIP-Chip; #820-male, ChIP-seq), we identified a common set of 708 genes which, with 1 kb flanking sequences, overlap H3K27me3 enriched chromatin regions. We queried these 708 putatively H3K27me3 regulated genes for expression in adult heads using the modENCODE high-throughput RNA-seq data (92) and selected a set of 16 genes for ChIP-qPCR analysis, including genes that are not expressed (*ato*, *eve*, *gsb-n*, *Ubx*; Reads Per Kilobase of transcript per Million mapped reads (RPKM) = 0), or expressed in low (*eyg*, *caup*, *E5*, *dve*, *mirr*; RPKM < 10), moderate (*Idgf3*, *hth*, *mAcR*, *srp*, *nAcR-beta21C*; 20 < RPKM < 50) or high levels (*ninaC*, *Obp99c*; RPKM > 200) in heads of 4 day old females. We prepared chromatin from heads of 3 day old Httex1p-Q93 (mutant) or Httex1p-Q20 (control) expressing flies and performed ChIP-qPCR analysis with H3 and H3K27me3 specific antibodies (Fig. 5). The H3K27me3 signal was at least an order of magnitude higher at all of the selected loci than in a control genomic region, F22. Furthermore, we found a moderately negative correlation between gene expression (modENCODE RNA-seq) and H3K27me3 enrichment data in both flies expressing mutant (R=-0.58, P=0.019) or control (R=-0.47, P=0.067) HTT. Surprisingly, however, a significant change in H3K27me3 levels between the mutant and control could not be detected at any of the inspected genes. This, together with the immunoblot data suggest that H3K27me3 methylation balance is not disturbed in mutant HTT challenged flies, therefore the rescuing effect of Utx is likely not acting by restoration of wild-type H3K27 methylation patterns.

**Figure 5.**
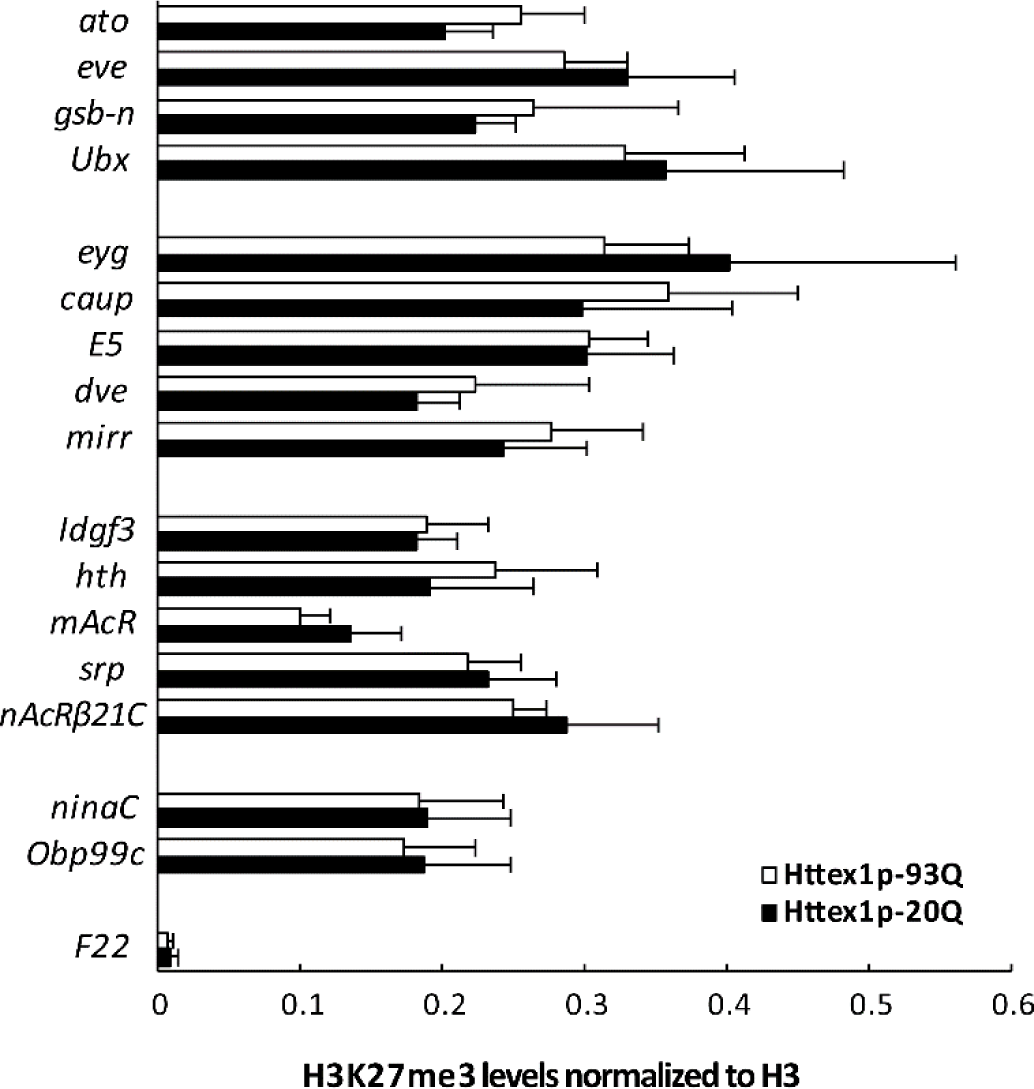
H3K27me3 occupancy is not altered on candidate genes in mutant Htt expressing flies. The level of H3K27me3 modification on genes with H3K27me3 enriched regions was determined by ChIP-qPCR in head samples of Httex1p-Q93 or Httex1p-Q20 (control) expressing flies. The tested gene set included genes not expressed in neurons (*ato*, *eve*, *gsb-n*, *Ubx*), and ones with low (*eyg*, *caup*, *E5*, *dve*, *mirr*), moderate (*Idgf3*, *hth*, *mAcR*, *srp*, *nAcR-beta21C*) or high (*ninaC*, *Obp99c*) expression levels. Compared to the F22 negative control region, H3K27me3 was enriched at all inspected genes but no significant change in H3K27me3 levels between Httex1p-Q93 and Httex1p-Q20 expressing flies was detected at any of the genes. Bars show the ratio of average ChIP-qPCR signals for H3K27me3 normalized to total input control and that for histone H3. Error bars represent standard error of mean (n=3).

### Direct manipulation of H3K27 residue by PTM mimetics

As the primary function of Utx is thought to be the demethylation of the H3K27 lysine residue, it was surprising that we could not detect alteration in trimethylated-H3K27 levels in HD flies. To further investigate whether the methylation state of the H3K27 influences HD pathology, we decided to manipulate this residue directly using post-translational modification mimicking transgenes. The non-essential endogenous His3.3A gene is located outside of the histone gene cluster, transcribed in a replication independent manner and was shown to replace canonical H3 in active genes (93, 94). We generated flies carrying transgenes of histone variant His3.3A (UAS-His3.3A) or its point mutant forms mimicking His3.3A being methylated (UAS-His3.3A-K27M, lysine mutated to methionine) or unmodified (UAS-His3.3-K27R, lysine mutated to arginine) at the K27 position. To test the effects of His3.3A-K27 PTM mimetics on HTT induced phenotypes while minimizing developmental defects induced by histone mimetics, we limited transgene expression to the adult nervous system using a heat sensitive allele of the GAL80 transcriptional regulator (GAL80^ts^) that inhibits GAL4 driven transcription at 18 °C but relieves it from repression at 30 °C. To reduce *His3.3A* gene load, we performed these experiments on flies heterozygous or homozygous for the *His3.3A^KO^* null mutation. For the tests, we raised flies expressing UAS-Httex1p-Q120 and one of the UAS-His3.3A transgenes simultaneously under the combined influence of the elav-GAL4 driver and tub-GAL80^ts^ at 18 °C, and transferred them to 30 °C after they eclosed.

Since it is not practical to score for eclosion rate and rhabdomere changes in ommatidia in those flies, we tested other aspects of HD pathology induced by HTT expression, e.g. reduced longevity and impaired motor function (95). First, we determined the longevity of male HD flies expressing His3.3A PTM mimic mutations. In general, we found the shape of the survival curves of different experimental categories similar with 9-11 days median and 13-15 days maximum lifespans (Fig. 6A and 6C). The restricted mean lifespan of males heterozygous for *His3.3A^KO^* and overexpressing Httex1p-Q120 along with either His3.3, His3.3A-K27R or His3.3A-K27M was 9.85, 10.09, and 9.75 respectively (Fig. 6B). Based on statistical analysis of the data, the lifespan difference of HD flies overexpressing His3.3A-K27R and controls overexpressing His3.3A is statistically significant (P=0.04, log-rank test with Bonferroni correction), although the effect size is minute. The restricted mean lifespan of males homozygous for His3.3A^KO^ and overexpressing Httex1p-Q120 along with either His3.3, His3.3A-K27R or His3.3A-K27M was 10.62, 10.74, and 9.97 respectively (Fig. 6D). In this case the lifespan difference between HD flies overexpressing His3.3A-K27M, and His3.3A or His3.3A-K27R are both statistically significant (P=0.0009 and P=0.0001, respectively, log-rank test with Bonferroni correction). Thus, survival data suggest that introduction of His3.3A-K27 methylation mimetics slightly decrease survival while hindering modification of the K27 residue slightly increases it.

**Figure 6.**
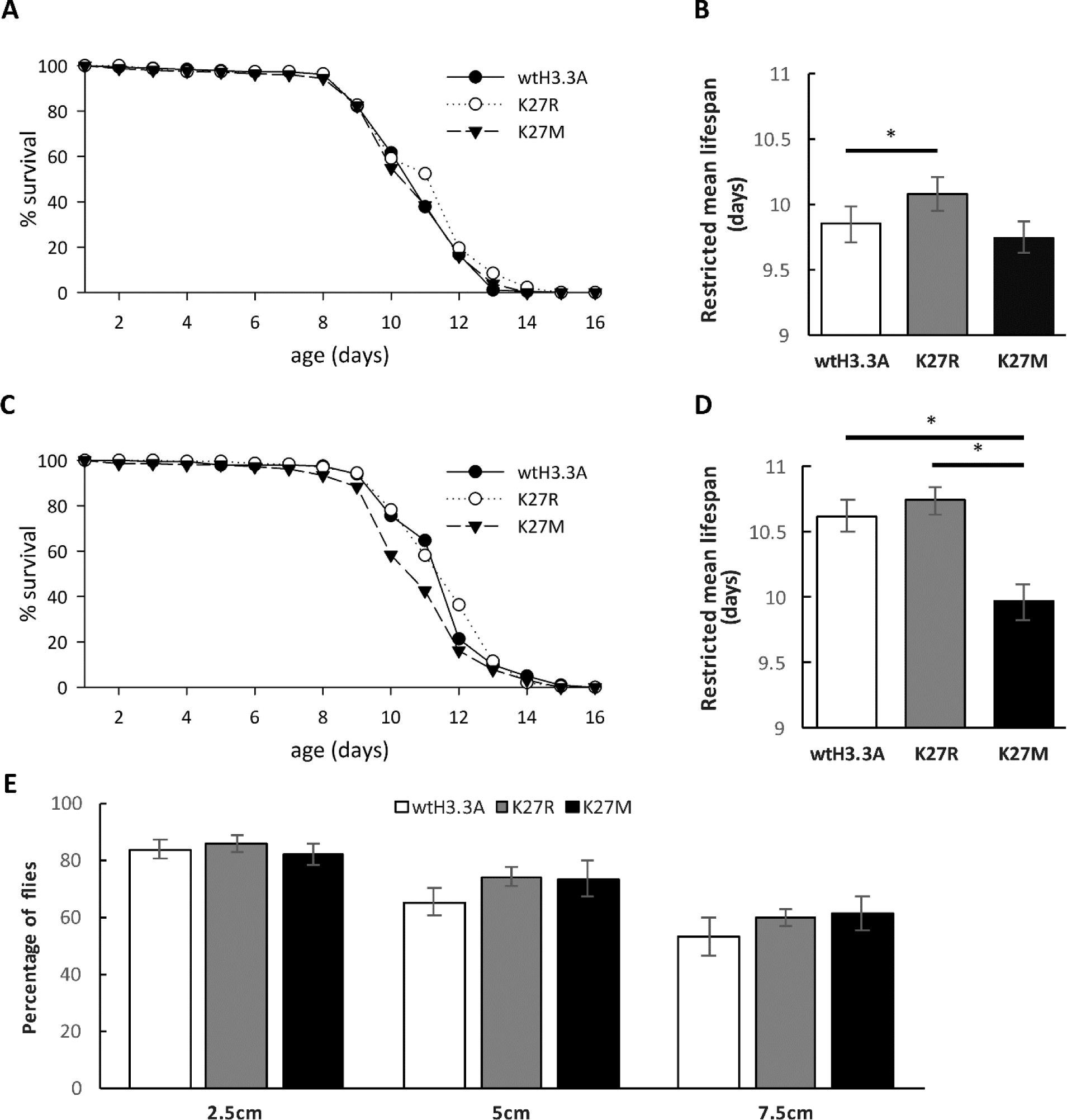
Direct manipulation of the H3K27 residue by PTM mimetic exert minor influence on HD pathology. The effects of H3K27 modifications on mutant Htt induced reduced longevity and impaired motor activity were investigated in flies co-expressing UAS-Httex1p-Q93 and wild-type or point mutant UAS-His3.3A (wtHis3.3A, His3.3A-K27R or His3.3A-K27M) in the nervous system in a hetero- or homozygous *His3. 3A^KO^* background. (**A**) Survival plot shows percent survival of Httex1p-Q93 expressing males heterozygous for *His3.3A^KO^* and co-expressing wtH3.3A (n=185), H3.3A-K27R (n=225) or H3.3A-K27M (n=255) as a function of time. (**B**) In heterozygous *His3.3A^KO^* background expression of His3.3A-K27R slightly but significantly increases longevity (P=0.0426, log-rank test) of HD flies. Bars show restricted mean lifespan. Error bars represent standard error of mean. (**C**) Percent survival of Httex1p-Q93 expressing males homozygous for *His3.3A^KO^* and co-expressing wtH3.3A (n=201), H3.3A-K27R (n=234) or H3.3A-K27M (n=216) as a function of time. (**D**) In homozygous *His3.3A^KO^* background expression of His3.3A-K27M significantly decreases longevity (P=0.0009, log-rank test) of HD flies. Restricted mean lifespan is shown. Error bars represent standard error of mean. (**E**) Motor activity of flies co-expressing UAS-Httex1p and wild-type or point mutant UAS-His3.3A were analyzed by the climbing assay. No significant difference between HD flies expressing wtH3.3A, H3.3A-K27R or H3.3A-K27M was found for the capability to climb at least 2.5, 5 or 7.5 cm vertically in 10 seconds.

Next, we used a climbing assay to determine the motor abilities of HD flies expressing histone PTM mimetics. To characterize climbing ability, we determined the percentage of 10 day old HD flies overexpressing His3.3A, His3.3A-K27R, or His3.3A-K27M on a homozygous *His3.3A^KO^* background that climbed at least 2.5, 5 or 7.5 cm vertically in 10 seconds (Fig. 6E). We found no significant differences in the climbing abilities of flies of different genotypes in these tests. Both His3.3A-K27R and His3.3A-K27M overexpressing flies performed slightly better than His3.3A control, but these differences were not significant statistically (P=0.29, ANOVA).

### Arginine methyltransferases

Beside lysine methyltransferases, we also tested three protein arginine methyltransferases (PRMT) that fall into the Type I and Type II PRMTs protein groups. Members of both groups are capable of both monomethylating and dimethylating arginine. However, while Type I enzymes dimethylate the same terminal N atom of the guanidinium group of the Arg side chain generating asymmetric dimethylarginine (aDMA); Type II enzymes catalyze the addition of a methyl group to each of the terminal N atoms, generating symmetric dimethylarginine (sDMA) (96).

Art3 is the ortholog of human PRMT3 whose targets are unknown. We tested two mutant alleles: *Art3^MI03542^*, in which the second exon is disrupted by a Minos element, and *Art3^c04615^*, a PiggyBac transposon insertion in the first intron resulting in the reduction of the mRNA level of the gene below 1% of control (q-RT-PCR data, not shown). Heterozygosity of *Art3^MI03542^* significantly increased the viability of HD flies (1.47 eclosion rate, P=0.0005, Fig. 7A) but did not have a significant effect on neurodegeneration (+0.34 rhabdomeres per ommatidium in *Htt Art3^MI03542^*/+ vs *Htt* +/+ control, N.S., Fig. 7B). The *Art3^c04615^* allele did not have a significant effect on either neurodegeneration (+0.35, N.S., Fig. 7B) or viability (0.77 eclosion rate, N.S., Fig. 7A).

**Figure 7.**
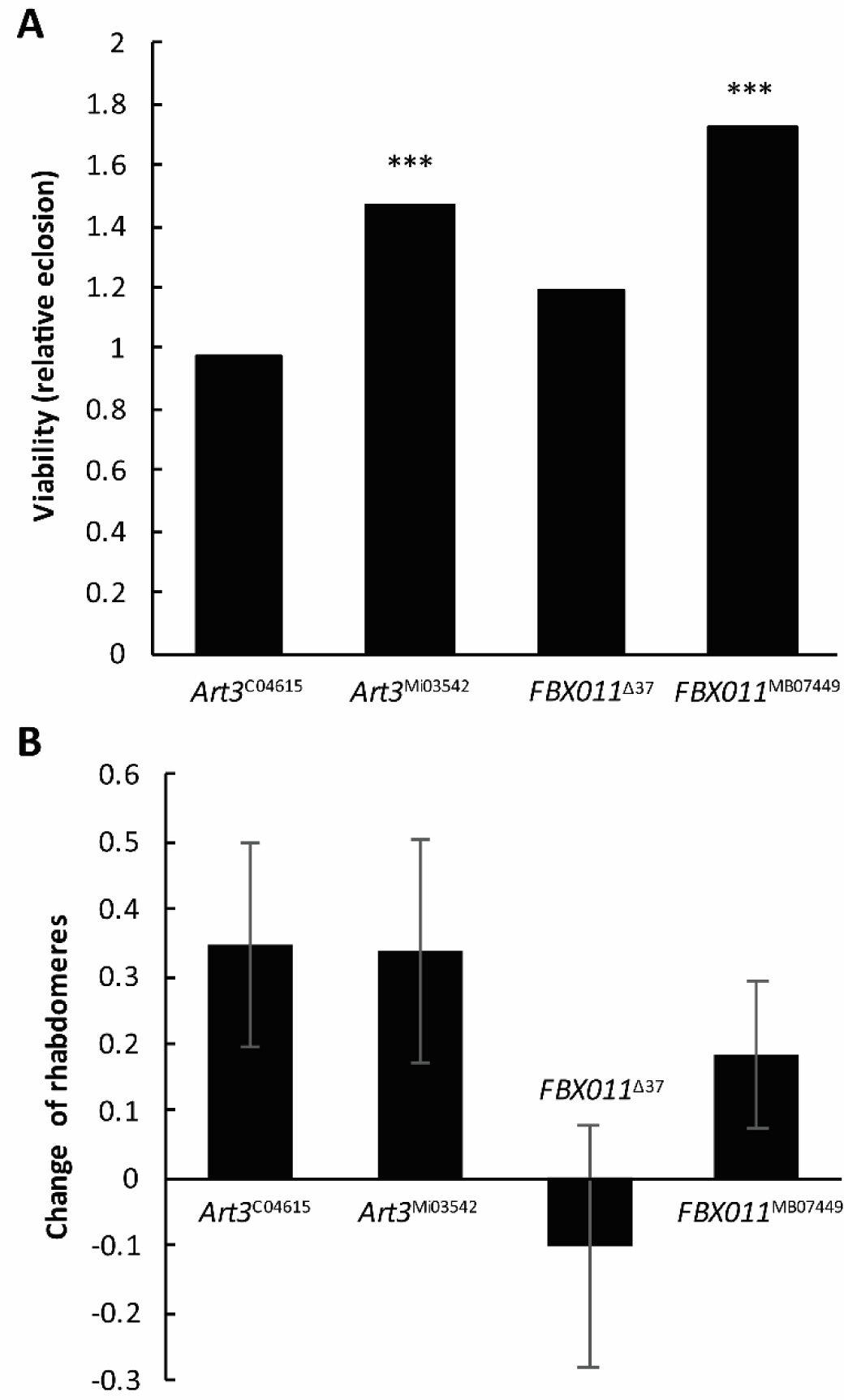
The effects of heterozygous reduction of protein arginine methyltransferases (PRMT), *Art3* or *FBXO11* on mutant Htt induced neurodegeneration and reduced viability. (**A**) Heterozygosity for the *Art3^MI03542^* or *FBX011^MB07449^* alleles significantly increased relative eclosion rate of Httex1p-Q93 expressing flies (P=0.0005 and P=4.5x10^-6^, respectively), while other alleles, *Art3^c0461^*and *FBX011^Δ37^*, did not have a significant effect. (**B**) None of the inspected alleles for *Art3* or *FBXO11* influenced the degeneration of photoreceptor neurons significantly. Bars show the difference of averages of number of rhabdomeres per ommatidium in the eyes of heterozygous PRMT mutant Htt expressing flies and Htt expressing control flies. Error bars represent standard error of mean.

*FBX011* (CG9461), the ortholog of human FBXO11/PRMT9, is a Type II methyltransferase that is also a component of an SCF ubiquitin ligase (E3) complex. FBX011 participates in the regulation of apoptosis (97) and miRNA and siRNA silencing (98). Heterozygosity of *FBX011^MB07449^* allele, in which a Minos element disrupts the first exon of the gene, caused increased viability of HD flies (1.72 eclosion rate, P=4.5×10^-6^) while that of *FBX011^Δ37^* deletion allele did not have significant effect on the viability (relative eclosion rate of 1.19, N.S., Fig. 7A). Neither of the *FBX011* alleles have influenced neurodegeneration (change of rhabdomeres per ommatidia is −0.10 in *Htt FBX011^Δ37^*/+ vs control and +0.19 for *Htt FBX011^MB07449^* vs control, N.S., Fig. 7B).

## DISCUSSION

Besides Huntington’s disease, pathology in a host of other CNS disorders has been correlated with transcriptional dysregulation including other polyQ diseases (99, 100), Alzheimer’s (101), Parkinson’s (102), Amyotrophic Lateral Sclerosis (ALS) (101) and Rubinstein-Taybi syndrome (103). In this study we address the question whether enzymes affecting the status of protein lysine methylation, a posttranslational modification that has a profound effect on chromatin structure and gene expression, influence the pathology of HD. By altering the level of enzymes responsible for the correct formation of lysine methylation patterns, we found that methyltransferases and demethylases affect HD pathology in an intricate manner that suggests target locus specific effects and raises the issue of methylation of non-histone proteins.

Our data reveal that manipulation of enzymes that regulate histone marks characteristic of constitutive heterochromatin has little or no effect on HD pathology. We found that methyltransferases modifying the H4K20 residue, PR-Set7 and Hmt4-20, did not exert a significant effect on HD pathology. Among the five H3K9 specific factors tested, only two influenced Htt induced phenotypes: reducing Su(var)3-9 enhanced while reducing JHDM2 suppressed the negative effect of mutant huntingtin on viability. However, neither of these factors had an impact on degeneration of photoreceptor neurons. These results show that manipulation of several categories of methyl modifying loci associated with heterochromatin formation does not affect HD pathology and suggest that transcriptional dysregulation in HD is not due to major reconfiguration of constitutive heterochromatin. Thus, the enzymes acting on constitutive heterochromatic marks can be eliminated as attractive therapeutic candidates.

Analysis of functionally related genes encoding enzymes modifying activating epigenetic methyl marks –including methyltransferases and demethylases acting on the H3K4 residue, which is trimethylated on promoters of active genes, or on the H3K36 or H3K79 residues that are methylated on the body of transcribed genes – provided ambiguous results. We have previously reported reduced H3K4 trimethylation on promoters of down-regulated genes in R6/2 mice and human HD samples and showed that partial loss of the H3K4 trimethyl specific demethylase Lid ameliorated HD pathology in *Drosophila* (7). The altered state of H3K4 methylation in post-mortem brain samples of HD patients was also observed in a study conducted by Dong and coworkers (104). However that analysis found no correlation between H3K4me3 levels at promoters and transcription levels, thus raising the possible involvement of non-histone targets of these enzymes or more distal sites of action at distant enhancers of prefrontal cortex genes.

In this study we found that reducing other H3K4 specific demethylases beside Lid, i.e. Su(var)3-3 and Kdm2, also rescued mutant Htt induced toxicity and neurodegeneration. Unexpectedly, we found that reduction of two out of the three tested H3K4 specific methyltransferases, Trx and Trr also suppressed neurodegeneration, although their effect on viability was opposite. Experiments aimed at testing enzymes modifying the H3K36 or H3K79 residues gave similarly gene specific results: while partial loss of the H3K36 specific methyltransferase Set2 ameliorated neurodegeneration, reduction of H3K36 specific Ash1 and the H3K79 specific methyltransferase Gpp enhanced it. Furthermore, heterozygous loss of the H3K36 specific demethylase Kdm4A had no effect on degeneration of photoreceptor neurons but increased viability.

The fact that modulating both the methylating and demethylating activities can have the same effect on HD pathology (e.g. H3K4 methylases and demethylases) points to the possibility of a complex set of epigenetically regulated disease modifying target loci and/or non-histone protein targets for those methylation modifying enzymes, which makes them a challenging therapeutic target.

One of the most important findings of this study is that manipulation of enzymes that affect facultative heterochromatin (which is typically dynamically regulated chromatin) impact HD pathology. Specifically, we found that loss of function mutations of the catalytic subunit of the PRC2 complex, E(z), that is responsible for H3K27 trimethylation, enhances neurodegeneration, and similar effects were observed with other components of the PRC2 complex, Esc and Su(z)12. In contrast, reducing the amount or activity of the H3K27 specific demethylase, Utx, by genetic or pharmacological means ameliorated neurodegeneration and reduced HTT aggregation. These findings that identify Utx as an appealing target for potential HD therapeutic intervention and are consistent with previous studies demonstrating the involvement of misregulated H3K27 methylation in several neurodegenerative disorders. For example, the murine PRC2 complex was shown to have a critical role in the maintenance of proper gene expression in medium spiny neurons (MSN) that are most affected in HD, by repressing non-neuronal, non-MSN specific and cell death promoting genes (105). PRC2 deficiency of *Ezh1* and *Ezh2* in MSN neurons resulted in progressive fatal neurodegeneration, reduction of the number of striatal neurons and impaired motor performance with concurrent upregulation of non-MSN-expressed genes and downregulation of MSN-specific genes (105). In contrast, in the neurodegenerative disease ataxia-telangiectasia (A-T), reduced activity of PRC2 suppresses pathology. In a murine *Atm*^-/-^ knock-out model of A-T, increased stability and protein level of EZH2 led to a substantial increase in the number of H3K27me3 binding sites on chromatin and concurrent reduction of gene expression. This transcriptional change had a significant impact on A-T pathology as knock-down of *Ezh2* in *Atm*^-/-^ mice was protective, resulting in reduced caspase-3 activation and Purkinje cell degeneration, and increased motor performance (106). PRC2 also modulates the severity of Spinal Muscular Atrophy (SMA) by regulating the activity of the *Survival Motor Neuron 2* (*SMN2*) gene, which is a duplicated gene highly similar to *SMN1* that is mutated in SMA (107). In the case of Huntington's disease, beside the potential effects of PRC2 activity on neurodegeneration, the role of normal huntingtin in the regulation of protein methylation by PRC2 might influence pathology. It was previously reported that full-length murine huntingtin physically interacts with Ezh2 and Suz12 and increases the methyltransferase activity of PRC2 *in vitro* (80). Accordingly, mouse embryos lacking wild-type *huntingtin* exhibited decreased trimethylation of histone H3 at lysine 27 and failed to repress PRC2 regulated Hox gene expression (80).

Despite the phenotypic effects of the opposing H3K27 specific methyltransferase and demethlyase in our model, we did not find direct physical evidence that would suggest direct mHtt induced changes in histone H3K27 modification: altered H3K27me3 levels could not be detected by either immunoblot or ChIP assays, and direct genetic manipulation of the H3K27 residue did not replicate the phenotypic changes observed in *E(z)* or *Utx* mutants. This raises the possibility that improved pathology in response to manipulation of methyl modifying enzymes may be due either to non-histone targets of these proteins or to putative demethylase independent activities.

The activities of several methyltransferases and demethylases are not restricted to histones but they can also modify other proteins. For example, EZH2 methylates the transcriptional factors STAT3 and PLZF, which in the case of STAT3 leads to increased phosphorylation and enhanced activity (108) while in the case of PLZF, it leads to ubiquitylation and degradation of PLZF (109).

Another possibility besides the potential targeting of non-histone proteins is that the effects that Utx mutants exert on HTT toxicity may be independent of the demethylase activity of the enzyme. It was demonstrated previously that mouse UTY, a Y chromosomal catalytically inactive paralog of UTX, partially complements the effects of loss of UTX activity (110). While *Utx^-^* females and *Utx^-^*/*Uty^-^* males never pass embryonic stage E18.5 of development, a significant portion of hemizygous *Utx* null males with a wild-type copy of *Uty* reach adulthood, suggesting that demethylase independent activities of UTX/UTY might be crucial during development. These activities might still involve transcriptional regulation, as both UTX and UTY were shown to directly bind to and potentiate the activity of heart specific transcriptional factors (110, 111). Furthermore, UTX and UTY both physically interact with RBBP5, an H3K4 methyltransferase complex subunit, and the SWI/SNF chromatin remodeler BRG1 (110, 112). *Drosophila* UTX is involved in similar protein interactions as its mammalian counterpart including direct binding to the BRG1 homolog nucleosome remodeler BRM (113) and being a subunit of the TRR containing COMPASS-like H3K4 methyltransferase complex (33). These conserved interactions suggest that demethylation independent activities of *Drosophila* Utx might be also conserved. Thus, while transcriptional dysregulation is strongly correlated with HD and other neurodegenerative diseases and attention has focused on histone modifications, many of the enzymes that regulate these processes have non-histone targets and at least some of them also have additional catalytic activities that may contribute to the disease process. Similar to our previous study that pointed to HDAC1 and SirT1 and SirT2 (114), this study points to the PRC2 component modifying enzyme, Utx, as the most attractive therapeutic target among the *Drosophila* methyl modifying enzymes.

## MATERIALS AND METHODS

### Drosophila crosses

Flies were reared on standard cornmeal molasses medium at 25 °C unless noted otherwise. Genetic interaction crosses were done by first mating elav-GAL4; *Sp*/CyO; +/+ (in case of second chromosomal mutation) or elav-GAL4; +/+; *Sb*/TM6 (in case of third chromosomal mutation) females with males carrying the mutation or RNAi construct of interest; then crossing the elav-GAL4; mutation/*Sp*; +/+ or elav-GAL4; +/+; mutation/*Sb* F1 male progeny to *w*; +/+; UAS-Httex1p-Q93 females. All F2 females express the UAS-Httex1p-Q93 transgene under the control of the neuronal elav-GAL4 driver; half of these females carry the mutation of interest in a heterozygous form, in F2 males elav-GAL4 is absent therefore they do not express UAS-Httex1p-Q93. To test mutations located on the 1^st^ chromosome, we set up control crosses with *w^1118^* side by side with the experimental crosses in order to compare the viability and pseudopupil phenotypes between the mutation-carrying and control flies. Relative eclosion rates from ≥1000 segregants were calculated as the ratio of the number of Httex1p-Q93 expressing flies carrying the mutation of interest (elav-GAL4/+; *Htt*/+; mutation/+) and the number of Httex1p-Q93 expressing mutation-free control flies (elav-GAL4/+; *Htt*/+; +/+) divided by the ratio of the number of Htt non-expressing siblings carrying the mutation of interest (+/Y; *Htt*/+; mutation/+) and the number of Htt non-expressing mutation-free siblings (+/Y; *Htt*/+; +/+). P values were based on Fisher’s exact tests.

### Pseudopupil analysis

The insect compound eye is comprised of ~1000 ommatidia (individual eyes) each of which contains 7 visible photoreceptor neurons that can be visualized by the specialized light gathering cellular structure called a rhabdomere. Pseudopupil analysis allows one to count the number of surviving photoreceptor neurons by counting the light gathering rhabdomeres. Analyses were carried out on 7 day old flies, which were raised 22.5 °C and shifted to 25 °C upon eclosion as described in detail (115). Change of rhabdomeres in each ommatidium was calculated as the difference of the number of rhabdomeres between the Httex1p-Q93 expressing flies with or without the heterozygous mutation of interest and the error bars represent the standard error of the mean (SEM) of difference of means. Pseudopupils of ≥ 5 flies were scored for each genotype.

### Drug test

Crosses of elav-GAL4 (females) x Httex1p-Q93 (males) were carried out in cages at 23 °C. About 60-100 eggs collected overnight were transferred to vials of standard cornmeal molasses medium containing indicated concentrations of GSK-J4 as described (90) (Tocris Bioscience), 0.1% dimethyl sulfoxide (DMSO, Sigma-Aldrich, negative control) or 100 mM butyrate (BA, Sigma-Aldrich, positive control). Freshly eclosed virgin females were collected and passed in vials with the corresponding drug/control food every day and aged at 25 °C for 7 days for pseudopupil analysis as described (115).

### Generation of His3.3A transgenic flies

Genomic DNA was first prepared from wild-type flies with NucleoSpin Tissue kit (Macherey-Nagel) and the genomic region of His3.3A was amplified by PCR using Q5 high-fidelity DNA polymerase (New England Biolabs) with primers located upstream (His3.3A_gF) or downstream (His3.3A_gR) of the gene. The 2174 bp amplicon was inserted to pJET1.2 vector with CloneJET PCR cloning kit (Thermo Scientific). This clone was used as template in PCR to amplify *His3.3A* from the start codon to the last codon before the stop codon using His3.3A_E3C_F (having a KpnI site and an AAA Kozak sequence before the start codon) and His3.3A_E3C_R primers (having an EcoRI site). The resulting amplicon was cut with FastDigest KpnI and EcoRI enzymes (Thermo Scientific) and cloned to the corresponding sites of pENTRY3C Gateway entry vector. Site directed mutagenesis was performed by amplifying the whole pENTRY3C-His3.3A clone in a 22 cycle inverse PCR reaction with Q5 polymerase using a common forward primer (K27_F, see Table 1S for all the primer sequences used in the study) and reverse primers having base substitutions at the desired positions (K27M_R and K27R_R). After PCR amplicons were phosphorylated with T4 polynucleotide kinase (Thermo Scientific), circularized with T4 DNA ligase (Thermo Scientific), and treated with DpnI to degrade methylated template DNA. Modified and wild-type H3.3A inserts were subcloned to pTWF-attB Gateway destination vector that was modified to carry a Φ C31 attB sequence at its NsiI restriction site, then these constructs were injected to embryos carrying the *attP*-*zh86Fb* Φ C31 docking site to generate transgenic flies by site specific transgene integration. To validate transgenic strains genomic DNA was prepared from transgenic flies, transgene sequences were PCR amplified using pTWFattB_Fseq and pTWFattB_Rseq primers, and subjected to Sanger sequencing.

### Longevity and motor performance of His3.3 mutants

The effect of His3.3A point mutants on longevity and motor performance in the HD model was tested on elav-GAL4; *His3.3A^KO^* UAS-Httex1p-Q120/*His3.3A^KO^*; UAS-His3.3A/tub-GAL80^ts^ flies, in which the UAS-His3.3A transgene was either wild-type (UAS-His3.3A-wt), or had a lysine to arginine (UAS-His3.3A-K27R) or lysine to methionine (UAS-His3.3A-K27M) substitution. These progeny were generated by mating elav-GAL4; *His3.3A^KO^*; tub-GAL80^ts^ females to w; *H3.3A^KO^* UAS-Httex1p-Q120; UAS-H3.3A males, raised at 18 °C and transferred to 30 °C after eclosion to induce transgene expression in the adult nervous system. To determine longevity, flies were kept at 30 °C, transferred to fresh vials every second day and the number of deceased individuals recorded daily. At least 185 flies per genotype was used for longevity analysis.

Climbing assays on elav-GAL4/+; *His3.3A^KO^* UAS-Httex1p-Q120/*His3.3A^KO^*; UAS-His3.3A/tub-GAL80^ts^ flies were performed on males that were reared at 18 °C and transferred to 30 °C after eclosion. Groups of 10 days old flies were transferred to a glass cylinder and the distance climbed vertically in 10 seconds after knocked to the bottom was recorded on video for 8 trials per each group. At least 35 flies divided to several groups were tested for each genotype.

### Quantitative RT-PCR

Heads of elav-GAL4>UAS-Httex1p-Q93 with or without mutation (control) female flies were homogenized in TRIzol reagent (Invitrogen) and RNA was prepared according to the manufacturers’ recommendations. First strand cDNA was prepared from 1 μg of total RNA with Maxima Universal First Strand cDNA Synthesis Kit (Thermo Scientific) using random hexamer primers. The resulting cDNA was diluted 1:10 and quantified in qPCR reactions with gene specific primers (UAS-F and UAS-R for *Htt*, *G9a*-*F* and G9a-R for *G9a*, *Utx*-*F* and Utx-R for *Utx*) in an MJ Research Opticon thermal cycler using SYBR Green PCR Master Mix (Applied Biosystems). Transgene expression levels were determined compared to a template calibration curve and normalized to that of the *rp49* housekeeping gene.

### Immunostaining

Wandering third instar larvae reared at 25 °C were cut into half in a drop of 1X PBS and flipped inside out. After removing extra fat bodies and other tissues, the dissected larvae were incubated in fixative (4% paraformaldehyde and 0.2% triton X-100 in 1X PBS) for 20 minutes. The larvae were then washed and blocked with PBSTS (10% Bovine serum albumin in PBT) for 1 hour at RT. After that, the larvae were incubated in rabbit polyclonal HTT exon1-specific antibody (1:2000 VB3130, Viva Bioscience) overnight at 4 °C. Upon removal of the primary antibody, the larvae were blocked again with PBSTS for 1 hour and incubated with secondary antibody solution (1:10,000 Alexa 568) for 1.5 hours at RT. Finally, the eye discs were dissected and mounted in Vectashield antifade mounting medium (Vector Laboratories). Images were captured with a Zeiss 780 laser scanning confocal microscope (Carl Zeiss MicroImaging), using the accompanying Zeiss software. HTT aggregate-positive puncta were counted using the “3D Object Counter” plug-in in ImageJ on maximum-intensity complied 3D images. The threshold levels were set to 1800 where most of the aggregates can be clearly identified.

### Histone extraction and quantitative western analysis

For histone extraction, 3 day old female or male fly heads or L2 larvae were homogenized in 5% Triton buffer (1X PBS, 5% Triton X-100, 3 mM DTT, 1 mM sodium orthovanadate, 5 mM sodium fluoride, 1 mM PMSF, 5 mM sodium butyrate and 1 tablet of Roche’s Complete Mini Protease Inhibitor), spun down at 4,000g for 8 min at 4 °C and the pellets containing the nuclei were washed 2x times with the same buffer. The pellets were then resuspended in 0.2N HCl and sonicated for 2 times of 10 seconds with 1 min interval at 4 °C. The nuclear histone proteins were acid extracted for 3 hours at 4 °C. After centrifugation, the supernatant was neutralized and 2% deoxycholate was added to the final concentration of 125~250 μg/ml and the samples were incubated for 10 min at RT. Trichloroacetic acid was then added to a final concentration of 6% and the protein pellets were washed with cold acetone. The pellets were finally dried and dissolved in appropriate volume of 1X loading buffer (50 mM Tris, pH6.8, 2% SDS, 10% glycerol, 0.0015% Bromophenol Blue). The samples were boiled for 5 min at 95 °C and loaded to custom-made 15% Tris-Tricine polyacrylamide gel and run at a constant voltage of 120 V for about 1 hour until the dye reached the bottom of the gel. PVDF membrane was used for standard Western blotting.

For quantitative western analysis, blots were blocked for 1 hour at room temperature using 50% 1X PBS diluted Odyssey infrared imaging system blocking buffer (LiCor) and then incubated overnight at 4 °C with antibody against histone H3K27me3 (Active Motif # 39155, 1:1000). Following incubation with primary antibody, blots were washed and incubated with goat anti-rabbit IRDYE 680CW (Li-Cor) secondary antibody (1:20,000) for 1 hour at room temperature. Blots were washed and scanned using the Odyssey Infrared Imager (Li-Cor). After scanning, blots were incubated with the second primary antibody H2B (Active Motif #61037, 1:2000) overnight at 4 °C, then washed and incubated with goat anti-mouse IRDYE 800CW (Li-Cor) secondary antibodies (1:20,000), washed and scanned as above. Integrated signal intensity for histone modifications and total H2B was acquired using Odyssey Software. Data were exported to Excel and the ratios of signals from different histone modifications to total H2B obtained.

### Chromatin immunoprecipitation

Chromatin samples from heads of 3 days old *w* elav-GAL4/*w*; Httex1p-Q93/+ and *w* elav-GAL4/*w*; Httex1p-Q20/+ females were prepared as described previously (116) with modifications. In short, 1500-2500 fly heads were ground under liquid N2 and homogenized in homogenization buffer (15 mM HEPES pH 7.6, 10 mM KCl, 5 mM MgCl2, 0.5 mM EGTA, 0.1 mM EDTA, 0.1% Tween, 350 mM sucrose, with 1 mM DTT, 1 mM PMSF and Protease Inhibitor Cocktail (Merck)). Chromatin was crosslinked with 1% formaldehyde for 10 min at RT, then after collection and lysis of nuclei fragmented by sonication with a Bioruptor (Diagenode). Samples containing equal amounts of DNA as measured by a fluorometric method (Qubit 2.0 with BR DNA assay kit (Life Technologies)) were used for ChIP analysis. Diluted samples were pre-cleared with Dynabeads Protein A (Life Technologies) for 4 hours at 4 °C and then immunoprecipitated with 3 μg anti-H3 (Abcam ab1791) or anti-H3K27me3 (Millipore 07-449) ChIP-grade polyclonal antibody, or no antibody (mock control) at 4 °C overnight. Chromatin-antibody complexes were bound to Dynabeads Protein A for 4 hours at 4 °C. The supernatant of the mock control was collected and used as total input control (TIC). After extensive washes chromatin was reverse crosslinked overnight at 65 °C, treated with RNase A and Proteinase K, extracted with phenol-chloroform then precipitated with ethanol in the presence of glycogen. The amount of immunoprecipitated DNA was determined by quantitative PCR in a PikoReal Real-Time PCR system (Thermo Scientific) using Power SYBR Green Mastermix (Life Technologies). Gene specific primers used are listed in Supplementary Table 1. The quantity of immunoprecipitated DNA was determined by setting sample Ct values to a TIC calibration curve and deducting the amount of DNA in the mock controls.

## ACKNOWLEDGEMENTS

The authors wish to thank Drs. Jeff Simon for *E(z)^63^*, Andreas Bergmann for *Utx* alleles, Alex Mazo for *trr* alleles, Greg Shanower for *gpp* alleles and Konrad Basler for the *His3.3A^KO^* allele. We thank Dr. Judit Pallos for her insight and help at the beginning of the project. We thank the Bloomington Drosophila Stock Center (NIH P40OD018537), the Drosophila Genomics Resource Center (DGRC), Bloomington, IN, supported by NIH grant P40 OD010949 and the Vienna Drosophila Stock Center for providing fly stocks. The authors acknowledge the support of the Optical Biology Core facility and the Cancer Center Support grant of the University of California, Irvine (CC grant #CA62203).

